# DNA2 variant analysis supports the nuclease activity as a preferred therapeutic target

**DOI:** 10.64898/2025.12.14.693011

**Authors:** Katherine E. Baillie, Sijie Zhang, Veena Mathew, Troy Nations, Balakrishna Koneru, Peter C. Stirling

## Abstract

DNA2 is a combined nuclease and helicase that processes long 5’ flaps and other structures that occur during DNA replication and repair. Current models invoke an essential function of DNA2 in processing stalled replication forks to dampen toxic recombination based restart. As an essential replication stress response factor, DNA2 has been proposed as an anti-cancer drug target with several tool compounds developed. Here we sought to model inhibited DNA2 states using dominant genetics with separation-of-function alleles to identify the optimal mechanisms for DNA2 targeting. We find that expression of DNA2 nuclease-dead alleles exert dominant effects on fitness and DNA damage repair that are dependent on its helicase and RPA binding activities. Furthermore, we find an imperfect link wherein a subset of ALT+ models are hypersensitive to the presence of DNA2-nuclease dead protein. These results were replicated with a nuclease-specific inhibitor of DNA2. Together, our data suggests that DNA2 targeting in cancer should be nuclease focused and will rely on identification of specific biomarker subsets within ALT+ or other tumor cell states.

## Introduction

DNA2 is a combined nuclease-helicase with roles across genome maintenance and repair. It primarily acts as a 5’-3’ nuclease, digesting RPA-coated flaps of single-stranded DNA (Masuda-Sasa et al, 2006). In replication, DNA2 processes long flap structures on lagging-strand Okazaki fragments (Stewart et al, 2008; Levikova & Cejka, 2015). It has also been implicated in replication of difficult template structures at telomeres, centromeres and rDNA (Lin et al, 2013; Fernandez et al, 2025; Li et al, 2018). In repair, DNA2 couples with BLM helicase to generate long 3’ flaps at double-strand breaks for homologous recombination (Nimonkar et al, 2011). While DNA2 possesses an ATP-dependent helicase domain, it requires partnering with RecQ helicases for unwinding long tracts of DNA (Sturzenegger et al, 2014). At reversed replication forks, DNA2 processing with WRN helicase mediates fork restart (Thangavel et al, 2015). Current models suggest this role prevents inappropriate recombination and underlies the essentiality of DNA2 from yeast to humans (Duxin et al, 2012; Hudson et al, 2025). In human disease, partial loss-of-function mutations cause microcephalic primordial dwarfism or a Rothmund-Thomson like syndrome, illustrating the importance of normal DNA2 activity for development (Tarnauskaitė et al, 2019; Filho et al, 2023).

In cancer, DNA2 is frequently overexpressed across multiple tumor types (Peng et al, 2012; Folly-Kossi et al, 2023; Kumar et al, 2017). High DNA2 expression has been correlated to decreased survival in breast and serous ovarian cancers (Folly-Kossi et al, 2023; Peng et al, 2012). As a result of its retention in tumors and importance for genome maintenance, several tool compounds have been developed for DNA2 inhibition as prospective cancer therapies. C5 binds to an allosteric pocket which inhibits DNA binding, and consequently nuclease and helicase activity (Liu et al, 2016). Analysis of DNA fibers showed C5 treatment inhibited replication fork restart and sensitized cells to treatment with camptothecin (CPT) and PARP inhibitors. Treatment with d16, a C5 derivative, was also shown to synergize with cisplatin and PARP inhibitors, which was enhanced in cells expressing mutant P53 (Folly-Kossi et al, 2023). Subsequent work on compounds from the NSC library identified selective inhibitors of the DNA2 nuclease function, although with an unknown binding site (Kumar et al, 2017). Induction of Ras activation in pancreatic and breast cancer cell lines enhanced sensitivity to NSC-105808, highlighting the importance of DNA2 in oncogene activated replication stress. Findings from inhibitor studies demonstrate that there are cancer-specific contexts that enhance sensitivity of cells to DNA2 inhibition, driving its pursuit as a precision anti-cancer target.

More recently, roles for DNA2 in the alternative lengthening of telomeres (ALT) process have been proposed. ALT is a telomerase-independent mechanism for telomere extension found in 10-15% of cancers (Gaspar et al, 2018). ALT is most common in tumors of mesenchymal origin, including osteosarcoma and soft tissue sarcomas, as well as brain cancers such as astrocytoma and glioblastoma (Burrow et al, 2024; Heaphy et al, 2011). ALT is driven by telomeric replication stress induced by difficult templates such as non-coding TERRA R-loops or G-quadruplexes (G4s) (Yang et al, 2021; Yadav et al, 2022). Replication stress initiates recombination and Polδ mediated break-induced replication of the highly repetitive telomeric DNA (Roumelioti et al, 2016). Multiple mechanisms of ALT have been identified, including a RAD52-dependent and RAD52-independent pathway (Zhang et al, 2019). There is no unifying genetic alteration in ALT, although most ALT lines contain mutations in either ATRX or DAXX chromatin remodelers (Turkalo et al, 2023). ALT status is instead characterized by several biomarkers, including telomere length heterogeneity, C-circles, and ALT-associated PML (promyelocytic leukemia) bodies or APBs (Sohn et al, 2023; Hoang & O’Sullivan, 2020). APB formation involves phase separation of telomeric DNA, shelterin complex components, and DNA repair factors, held together by PML and mediated by SUMOylation (Henson et al, 2005). The proposed roles for DNA2 involve processing of difficult template structures at ALT. Jiang et al. (2024) found that DNA2 restrained replication stress at ALT through resection of long Okazaki fragment flaps generated by BLM helicase (Jiang et al, 2024). DNA2’s function in R-loop processing may also be significant as DNA2 depletion in ALT cells increases telomeric R-loops (Ragupathi et al, 2025). The same study identified a synthetic lethal interaction between DNA2 and FANCM in ALT cell lines.

There has been increasing interest in targeting DNA2 in a variety of cancer contexts. With both nuclease and helicase activity, and a wide variety of functions across the cell, questions remain on the most effective approach to targeting DNA2. Here we use separation-of-function mutants to model DNA2 inhibition. We identify nuclease dead alleles as dominant negative mutations which induce DNA damage and cellular arrest in ALT cell lines. This phenotype is dependent on DNA binding and helicase activity, suggesting a protein trapping mechanism reminiscent of successful chemotherapeutic mechanisms like those of topoisomerase or PARP inhibition (Murai et al, 2012; Pommier, 2013). Further we show that only a subset of ALT cell lines are sensitive to DNA2 nuclease mutation or inhibition. Our work provides an important nuance to the emergence of DNA2 as a target in ALT cancers: not all ALT states are equal, with only a subset responding to dominant negative DNA2. Future work to characterize robust biomarkers for sensitivity to potent and selective DNA2 nuclease inhibitors will advance DNA2 as a target in cancer therapy.

## RESULTS

### Nuclease dead mutations in DNA2 are dominant negative in U2OS cells

To identify dominant negative mutations in DNA2, we first looked to known mutations from human disease and biochemical studies (**Table 1, Fig. S1A**). Mutants were overexpressed in U2OS cells and examined by immunofluorescence for γH2AX foci formation (**Fig. S1B,C**). Only the nuclease dead alleles D277A and K300R increased γH2AX foci significantly above background and to the same extent as treatment with mitomycin C (MMC). These residues have roles in coordinating magnesium ions in the nuclease active site, indicating that the loss of nuclease catalysis is key to the observed phenotype (Zhou et al, 2015). In contrast, none of the alleles underlying human genetic diseases, nor the K654E helicase dead allele resulted in any DNA damage.

**Table 1.** Mutations in DNA2 and their functional consequences.

| Mutation | Domain | Functional result | Citations |
| --- | --- | --- | --- |
| D277A | Nuclease | Nuclease dead | (Masuda-Sasa et al, 2006; Nimonkar et al, 2011) |
| K300R | Nuclease | Nuclease dead in yeast DNA2 | (Balakrishnan et al, 2010) |
| T655A | Helicase | Microcephalic Primordial Dwarfism | (Tarnauskaitė et al, 2019) |
| K654E | Helicase | Helicase dead/ATPase dead | (Masuda-Sasa et al, 2006; Nimonkar et al, 2011) |
| R198H | Nuclease | Autosomal dominant progressive myopathy. | (Ronchi et al, 2013) |
| K227E | Nuclease |  |  |
| V637I | Helicase |  |  |

### Helicase activity is required for the nuclease dead phenotype

To assess the role of the helicase function in the dominant DNA damage phenotype, a double nuclease dead/helicase dead mutant was generated by site directed mutagenesis. The double mutant showed a complete rescue of the γH2AX phenotype (**Fig. 1A**). This indicates that the dominant negative mechanism of the nuclease dead mutant is dependent on helicase activity. To confirm that this result was not due to differences in protein expression, the expression of FLAG-tagged mutants was compared by western blot (**Fig. S2A**). The ND/HD mutant was expressed at the same or greater levels relative to the ND allele. The nuclease deficient DNA damage phenotype was also rescued in a K300R/K654E double mutant (**Fig. S2B**). Since the DNA2 helicase/ATPase is important for loading the enzyme onto long flaps, these data suggest that functional loading of the ND enzyme is essential to elicit the observed dominant induction of DNA damage. These results agree with previous experiments by Duxin et al. (2012), who demonstrated that expression of the D277A (ND) mutant could not be maintained in a transduced cell line (Duxin et al, 2012). However, a D277A/K654E (ND/HD) mutant deficient in both nuclease and helicase activity was tolerated.

**Figure 1.**
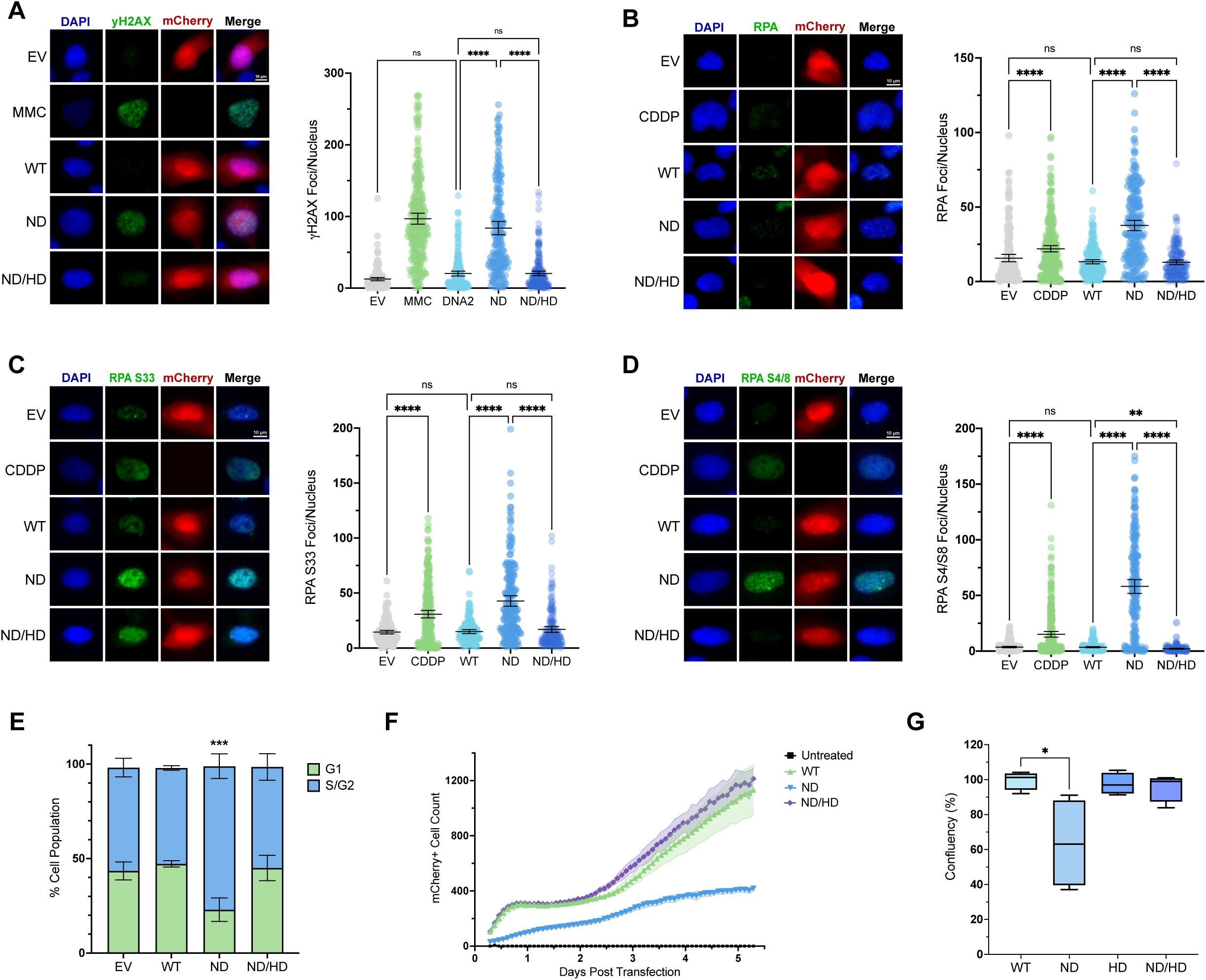
Expressing nuclease-deficient DNA2 causes dominant, helicase-dependent DNA damage and cell cycle delay. (A-D) Imaging and quantification of γH2AX, total RPA, phospho-Ser33-RPA32 and phospho-Ser4/8-RPA32 foci. For each panel, representative cell images are shown (left), with quantification and statistical comparison (right). Mitomycin C (MMC) or cisplatin (CDDP) treatments are used as positive controls, while empty vector (EV) or wildtype (WT) DNA2 transfection is a negative control. ND = nuclease dead, HD = helicase dead, ns = not significant. N=3; ****p<0.0001 by Kruskal-Wallis test; mean ± 95% CI. (E) Flow cytometry for DNA content following transfection of the indicated plasmids and quantification of G1 vs. S/G2 cells. ***P<0.001 by One-way ANOVA, comparing ND to all other conditions; mean ± SD. (F) Incucyte growth curve assay for U2OS cell proliferation over 5 days post-transfection with the indicated construct. The number of cells expressing the mCherry transfection marker was quantified by Incucyte for six technical replicates; mean ± 95% CI. (G) Viability of transfected cells was assessed 5 days post transfection by CellTitre-Glo® ATP assay. Minimum to maximum value shown with line at the median. N=4; *P<0.05 by t-test. For panels A-G, transfected cells were selected via a plasmid borne mCherry fluorescent marker.

### Nuclease dead DNA2 activates the replication stress response

As RPA-coated ssDNA is the endogenous substrate of DNA2, we next examined the impact of the ND mutant on RPA foci. Mutant expression increased RPA foci formation to a greater extent than treatment with cisplatin, a phenotype which was completely rescued in the ND/HD double mutant (**Fig. 1B**). This suggests that the ND mutant is not only preventing resection by endogenous DNA2, but impacting alternate mechanisms of 5’flap resolution leading to the accumulation of RPA foci. Accumulation of RPA-coated ssDNA triggers activation of the ATR complex (Zou & Elledge, 2003). In turn, activated ATR phosphorylates the RPA32 subunit on serine 33 (S33) (Maréchal & Zou, 2015). Rapid gathering of RPA at replication-associated DNA damage is associated with hyperphosphorylation of RPA32 (Liaw et al, 2011). Hyperphosphorylation can be detected as a second event at S4/S8, primarily mediated by DNA protein kinase (DNA-PK) (Liaw et al, 2011; Maréchal & Zou, 2015). The expression of the ND mutant significantly increased both the S33 and S4/S8 modifications of RPA (**Fig. 1C,D**). This indicates that the nuclease dead mutant is causing replication-associated DNA damage, activating replication stress and DNA damage response pathways.

Activation of the replication stress response would be expected to initiate cell cycle arrest pathways. Flow cytometry analysis of the cell cycle revealed accumulation in the S/G2 phase in cells expressing ND relative to WT or other mutants (**Fig. 1E**). To monitor the impact of DNA2 ND on cell growth, we used the ATP-glo assay and Incucyte monitoring, which both demonstrated significantly fewer ND cells relative to WT DNA2 after 5 days of transfection (**Fig. 1F, 1G**). These results coupled with the markers of stress response kinase activation suggest a G2/M checkpoint arrest in response to nuclease dead mutant expression.

### The nuclease dead mutant phenotype is mediated by interaction with DNA

It has been shown in several *in vitro* studies that nuclease dead DNA2 retains the ability to interact with 5’ flap ssDNA and displace RPA (Duxin et al, 2012; Pinto et al, 2016; Thangavel et al, 2015). This raises interesting possibilities as to the mechanism of the DNA2^ND^ dominant negative phenotype. For example, DNA2^ND^ may become retained on 5’ flaps without cleavage activity, in essence ‘trapping’ the protein. Another possibility is that the helicase activity of DNA2 processes and elongates the 5’ flap in an unregulated manner in the absence of nuclease activity. Both hypotheses require interaction with a DNA substrate. Therefore, we removed the RPA-interacting domains of DNA2 to test whether the phenotype is dependent on substrate binding. Versions of DNA2 lacking the α1 (DNA2^Δ1-20^) and α1-OB fold domains (DNA2^Δ1-121^) were generated, and expression was confirmed by western blot (**Fig. 2A,B**). Both deletions were then tested for induction of RPA phosphoserine 4/8 foci (**Fig. 2C**). The α1 deletion alone was not sufficient to rescue the phenotype, however deletion of the α1-OB fold domain resulted in a complete rescue. These results were supported by examination of γH2AX and RPA S4/S8 staining by flow cytometry (**Fig. S3**). The α1 domain interacts with external regions of RPA and is primarily involved in DNA2 recruitment (Zhou et al, 2015). In contrast, the OB domain forms interactions with RPA’s DNA binding domains, destabilizing the RPA-DNA attachment (Zhou et al, 2015). These results therefore suggest the phenotype requires the displacement of RPA on 5’ flap substrates by the ND mutant.

**Figure 2.**
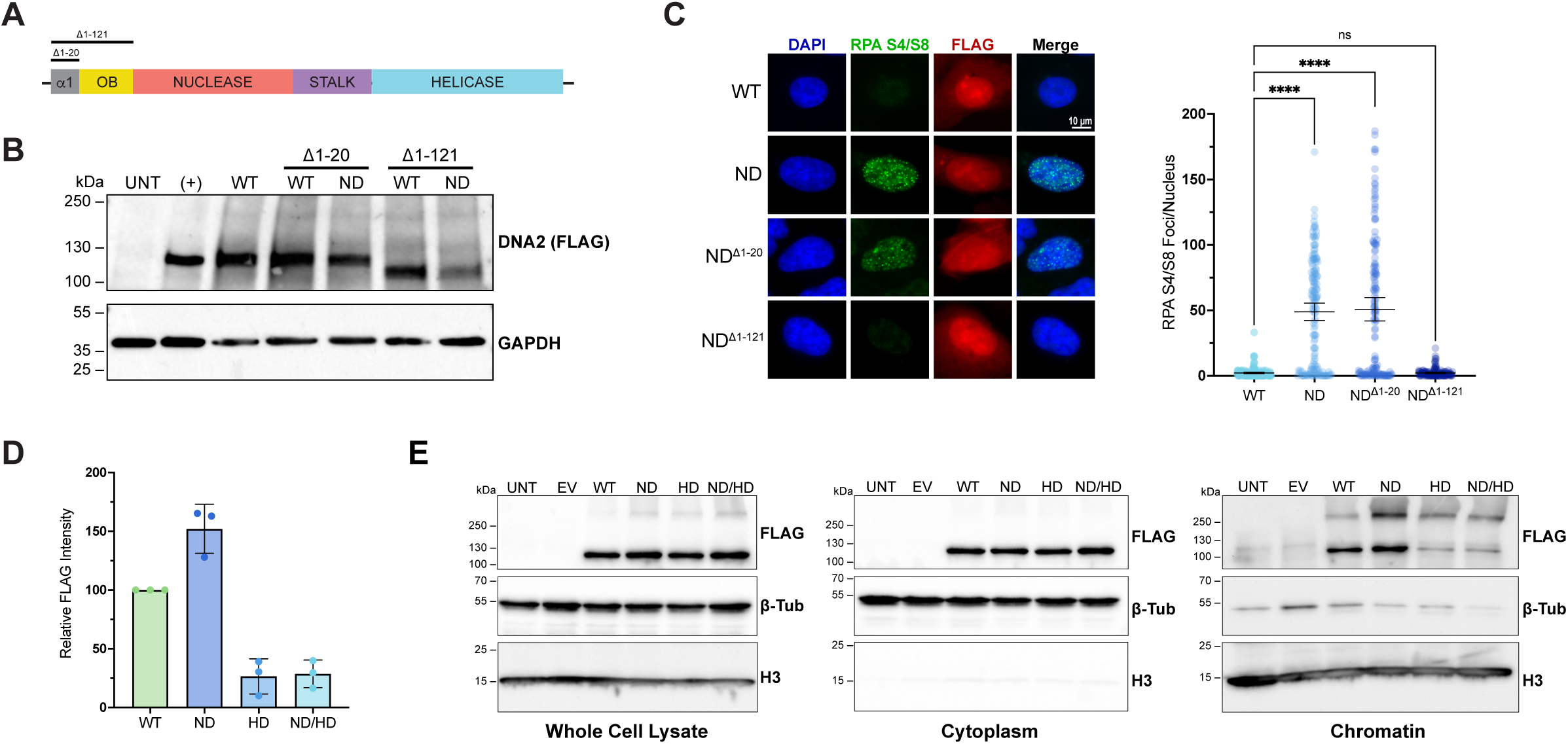
RPA displacement and chromatin loading drive dominant DNA2-ND phenotypes. (A) Schematic of DNA2 domains highlighting recruitment and displacement functions in the N-terminus. (B) Western blot of FLAG-DNA2 constructs and a GAPDH loading control. (C) Representative images of Phospho-Serine 4/8 RPA32 foci in the indicated cells. Here FLAG immunofluorescence marks transfected cells for foci quantification on the right. N=3; ****P < 0.0001 by Kruskal-Wallis test; mean ± 95% CI. (D) Relative quantification of FLAG-DNA2 in the chromatin compartment 48 hours post-transfection in HEK293T cells. Band intensity was normalized to total protein staining (Ponceau S) and plotted relative to WT sample for each replicate. N=3; mean ± SD. Helicase-dead alleles failed to accumulate normally on chromatin while ND was present in excess. (E) A representative chromatin fractionation result from the data quantified in D. β-tubulin (β-tub) and histone H3 (H3) are included as cytoplasmic and chromatin controls respectively.

We next used chromatin fractionation to study the interaction of ND DNA2 with DNA. Fractionation revealed increased ND in the chromatin compartment relative to wildtype, helicase dead, or double mutant (**Fig. 2D, 2E**). This result favours a ‘trapping’ mechanism whereby the nuclease dead mutant becomes loaded onto substrates that it cannot process, interfering with normal resection and DNA repair functions. Taken together, these experiments support a key role for interaction with RPA-coated DNA in mediating the dominant negative phenotype of ND DNA2 in cells.

### Cell lines with the ALT phenotype are sensitive to DNA2 nuclease dead mutants

Thus far, the dominant negative phenotypes of ND DNA2 have been primarily measured in the osteosarcoma derived U2OS cell line, which uses the ALT pathway for telomere elongation. Recent publications have identified roles for DNA2 in restraining replication stress at ALT telomeres through processing of long flaps created by BLM helicase and TERRA R-loops (Jiang et al, 2024; Ragupathi et al, 2025). To study the sensitivity of ALT and TERT cell line models to the ND mutant, we used flow cytometry staining of γH2AX and RPA phosphoserine 4/8 (**Fig. 3A**). Comparison of DNA damage following mutant expression revealed the VA-13 and SAOS2 ALT lines were similarly sensitive to ND expression, whereas HeLa, RPE1-hTERT and HeLa 1.3 cells were unaffected (**Fig. 3B**). The HeLa 1.3 cell line has exceptionally long telomeres, indicating that this effect was not due to telomere length or changes to telomere replication outside of the ALT context (Takai et al, 2010). However, the HEK293T cell line which is TERT+ showed similar sensitivity to U2OS (**Fig. 3B**). This may be partially influenced by higher levels of protein expression and agrees with our previous findings showing retention of DNA2^ND^ on chromatin in HEK293T cells (**Fig. 2E**). Nonetheless, the possibility of a connection between ALT status and the DNA2^ND^ phenotype is intriguing given the potential roles for DNA2 in these cancers. To further explore the association between telomere replication and the DNA2^ND^ phenotype, we used TeloFISH staining to study site-specific DNA damage. While there was an increase in RPA S33 foci co-localizing at TeloFISH signals (**Fig. 3C**), there is also a large population of S33 foci occurring outside of the telomeric regions. Therefore mutant DNA2 may be recruited to multiple sites in the genome such as reversed replication forks and long Okazaki fragment flaps to create damage.

**Figure 3.**
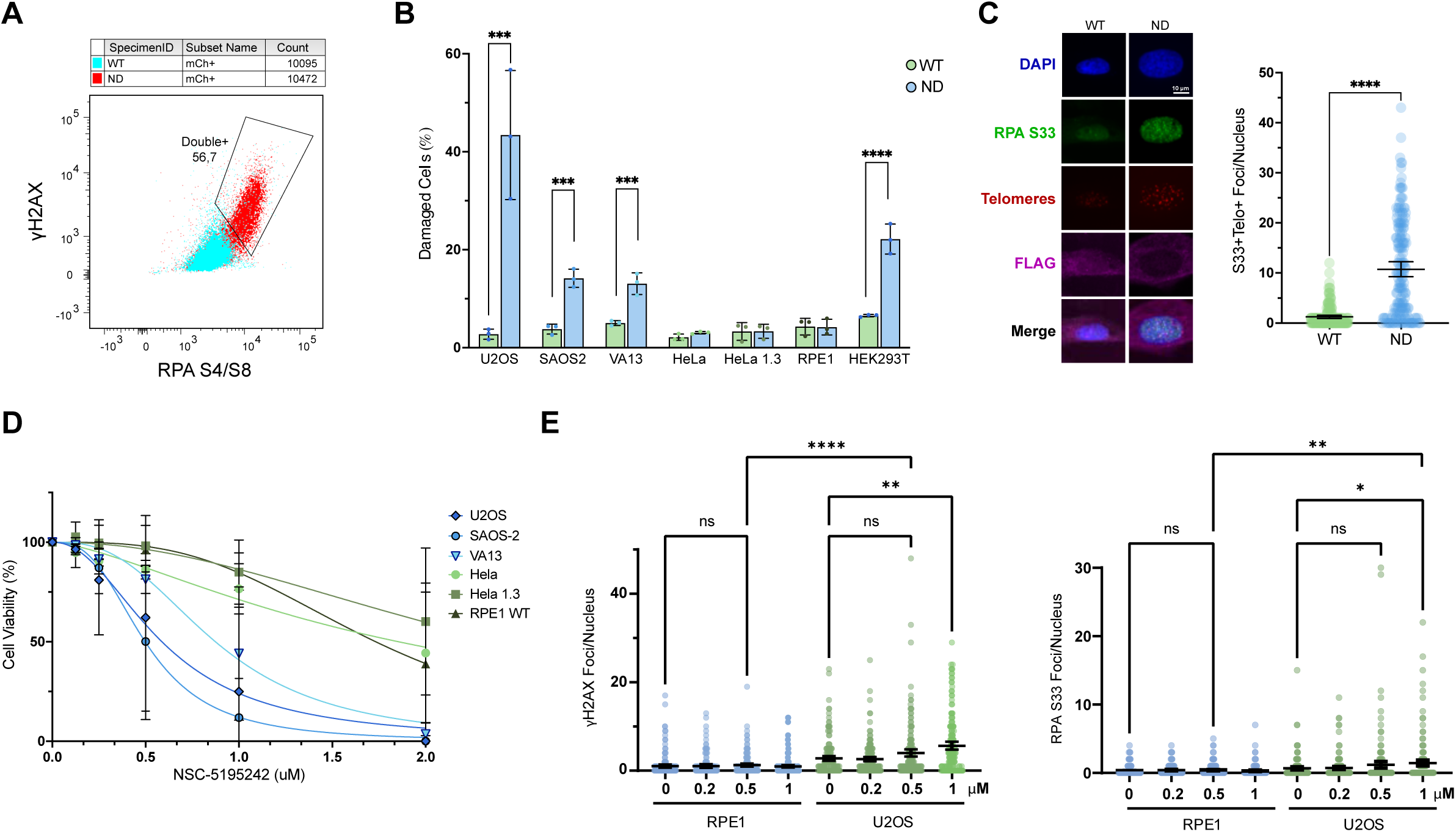
Telomeric damage and cell line selectivity of DNA2 nuclease inhibition. (A) Flow cytometry analysis of DNA damage (γH2AX) and replication stress (RPA ser4/8) markers in transfected cells (blue WT DNA2; red DNA2 ND). (B) Proportion of double positive cells by the flow assay as shown in A in the indicated cell lines with either WT-DNA2 or DNA2-ND expression. N=3; ***P < 0.001, ****P < 0.0001 by One-way ANOVA; mean ± SD.. (C) Representative images (left) and quantification (right) of co-localized telomeres and RPA phosphoserine 33 foci 48 hours post transfection in U2OS cells with the indicated DNA2 mutants. N=3; ****P < 0.0001 by Mann Whitney test, mean ± 95% CI. (D) Dose-response nonlinear regression curves of the indicated cell lines after treatment with serial dilutions of NSC-5195242. Cell viability quantified by CellTitre-Glo® ATP assay. ALT cell lines shown in blue, TERT+ in green. N=3; mean ± SD. (E) γH2AX (left panel) and RPA phosphoserine 33 (right panel) foci per nucleus of RPE1-hTERT or U2OS cells treated with the DNA2 inhibitor NSC-5195242 in increasing concentrations. Significant changes in DNA damage foci/nucleus in U2OS cells are indicated as N=3 with three technical replicates in each; ****p<0.0001,**P < 0.01, *P<0.05 by Kruskal-Wallis test; mean ± 95% CI.

### A DNA2 nuclease inhibitor replicates the DNA2 ND mutant phenotype

To explore the translation of findings from the ND mutant to existing DNA2 inhibitors, we tested a known DNA2 nuclease inhibitor NSC-5195242 against the panel of ALT and TERT positive cell lines (Kumar et al, 2017). This inhibitor was chosen as it is selective for the nuclease activity of DNA2 and should therefore align with our dominant ND mutant (i.e. a model wherein a proportion of cellular DNA2 is nuclease-deficient). As predicted, ALT positive cells were more sensitive to this inhibitor (IC50 0.50-0.87 μM) than TERT+ cell lines (IC50 1.73-2.41 μM), suggesting that NSC-5195242 mimics DNA2^ND^ (**Fig. 3D**). To determine if this inhibitor similarly causes DNA damage, γH2AX and RPA S33 foci were measured by immunofluorescence staining. DNA damage foci were induced upon NSC-5195242 treatment in U2OS cells in a concentration dependent manner but not in RPE1-hTERT cells (**Fig. 3E and Fig. S4a**). For comparison, we tested the DNA2 inhibitor C5, which interferes with the ability of DNA2 to interact with DNA, and found that this inhibitor was more potent in RPE1 than U2OS, supporting a different mechanism of action associated with complete DNA2 loss of function (**Fig. S4b**). This finding has significant implications for development of future DNA2 therapies as it suggests that total inhibition of DNA2 (i.e. by C5) may be broadly toxic, while inhibition of DNA2 nuclease activity could provide a window for selective killing of some cancers.

### DNA2 nuclease inactivation is detrimental to a subset of osteosarcoma lines independent of ALT status

Encouraged by our results with workhorse ALT cell line models, we elected to extend our analysis to a panel of patient-derived osteosarcoma (OS) cell lines. OS acquires an ALT phenotype in approximately 50% of patients, supporting its use as an ALT vs. TERT tumor model. We obtained three ALT lines (OS052, OS384, OS526) and three TERT lines (OS152, OS833, OS865) for analysis. ALT status was verified by an APB assay (**Fig. S5A**). Surprisingly only one of the tested ALT lines, OS384, exhibited increased γH2AX and RPA S4/S8 foci when DNA2-ND was expressed (**Fig. 4A-C**). However, two additional non-ALT lines, OS152 and OS833, showed significantly increased DNA damage foci with DNA2-ND expression (**Fig. 4A-C**). These results were further replicated with the DNA2 inhibitor NSC-5195242, which showed the same pattern of sensitive and resistant cell lines (**Fig. 4D**). This supports the view that NSC-5195242 is on-target, which we confirmed by CETSA (**Fig. S5B**). However, this indicates that ALT alone is not likely to be a suitable biomarker for DNA2 nuclease inhibitors in patient derived models broadly. Indeed, the strong response to DNA2 inhibition in some TERT model lines suggests that alternative vulnerabilities are likely to exist. Future work aimed at delineating which features of the ALT state, or other genome maintenance defects may create DNA2 nuclease dependencies are ongoing. This will be a critical step to advance a precision strategy for DNA2 nuclease inhibitor development.

**Figure 4.**
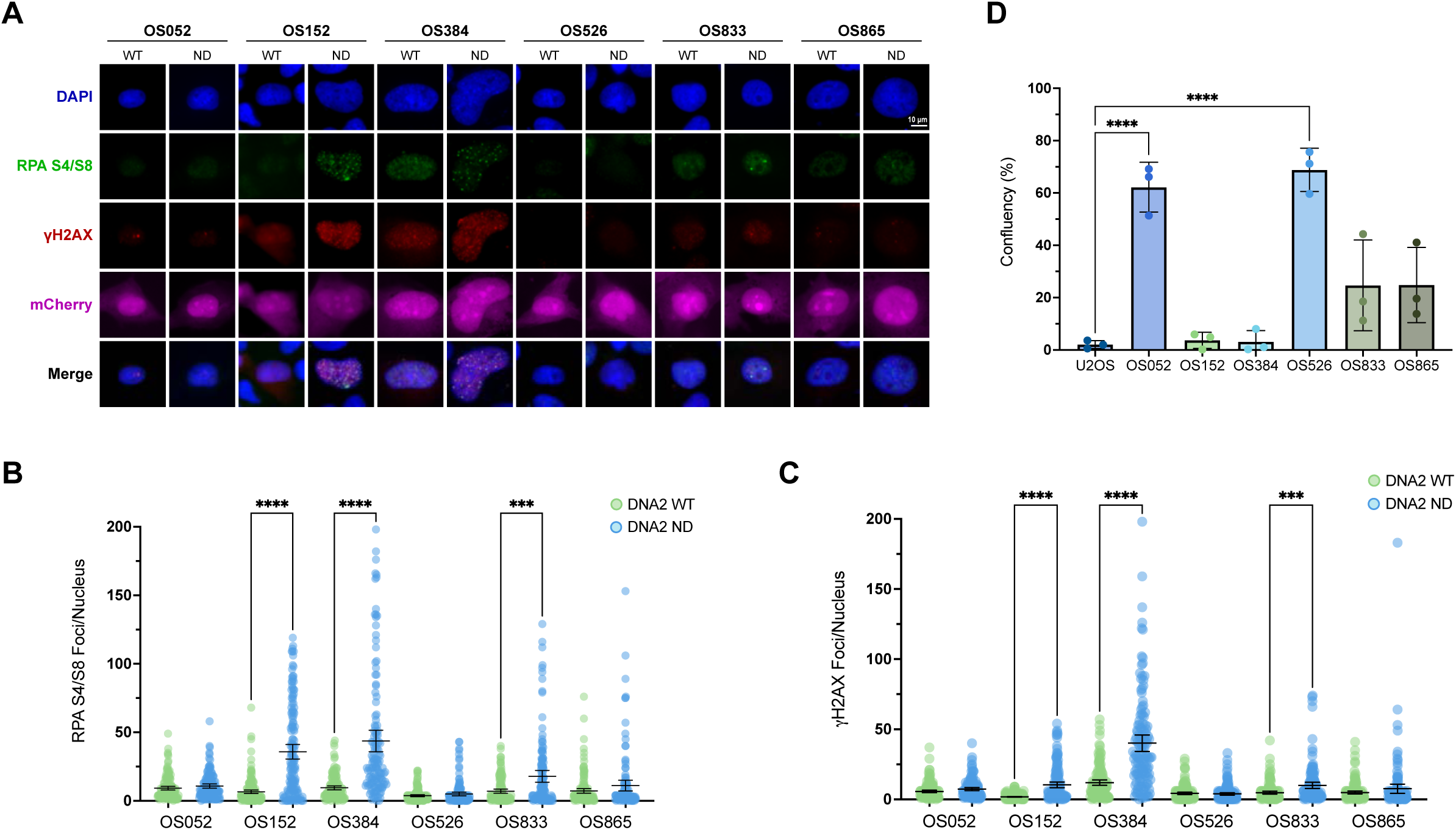
Differential sensitivity of patient derived osteosarcoma cell lines to DNA2 nuclease inhibition independent of ALT status. (A) Representative images of γH2AX and RPA ser4/8 staining in six OS cell lines with WT or ND DNA2 expression. (B and C) Quantification γH2AX and RPA ser4/8 foci from experiment in panel A. N=3; ****P<0.0001, ***P<0.001 determined by Kruskal-Wallis test; mean ± 95% CI (D) Cell survival of ALT (blue) and TERT (green) cell lines was measured 72 hours after treatment with 0.5 μM of the DNA2 inhibitor NSC-5195242. Cell viability quantified by CellTitre-Glo® ATP assay. N=3; ****P<0.0001 determined by ANOVA; mean ± SD.

### Replication fork reversal drives sensitivity to DNA2 nuclease inactivation

DNA2 has many potential substrates, including reversed replication forks, G-quadruplexes, R-loops, and long Okazaki fragment flaps. To understand which of these substrates may drive sensitivity to nuclease inactivation, we used siRNA knockdown of ZRANB3 to decrease replication fork reversal (**Fig. 5A**). Comparison of RPA S4/S8 foci induction upon ND expression between siCTRL and siZRANB3 showed a partial rescue of the phenotype (**Fig. 5A**). Western blot analysis of knockdown efficiency showed only partial knockdown, which may have contributed to this result (**Fig. 5B**). This suggests that an excess of reversed replication forks may be one substrate driving the dependency of cell lines to the DNA2 ND mutant. If true, inducing fork reversal in an insensitive cell line should sensitize to DNA2 nuclease inhibition. To test this, we treated RPE1-hTERT with hydroxyurea (HU) to stall replication, added a sub-lethal dose of NSC-5195242 and monitored DNA damage using γH2AX foci (**Fig. 5C**). While this concentration of NSC-5195242 had no effect in RPE-hTERT alone, when combined with HU it strongly induced DNA damage, supporting the view that stalled replication intermediates could be targeted by DNA2 nuclease inhibition.

**Figure 5.**
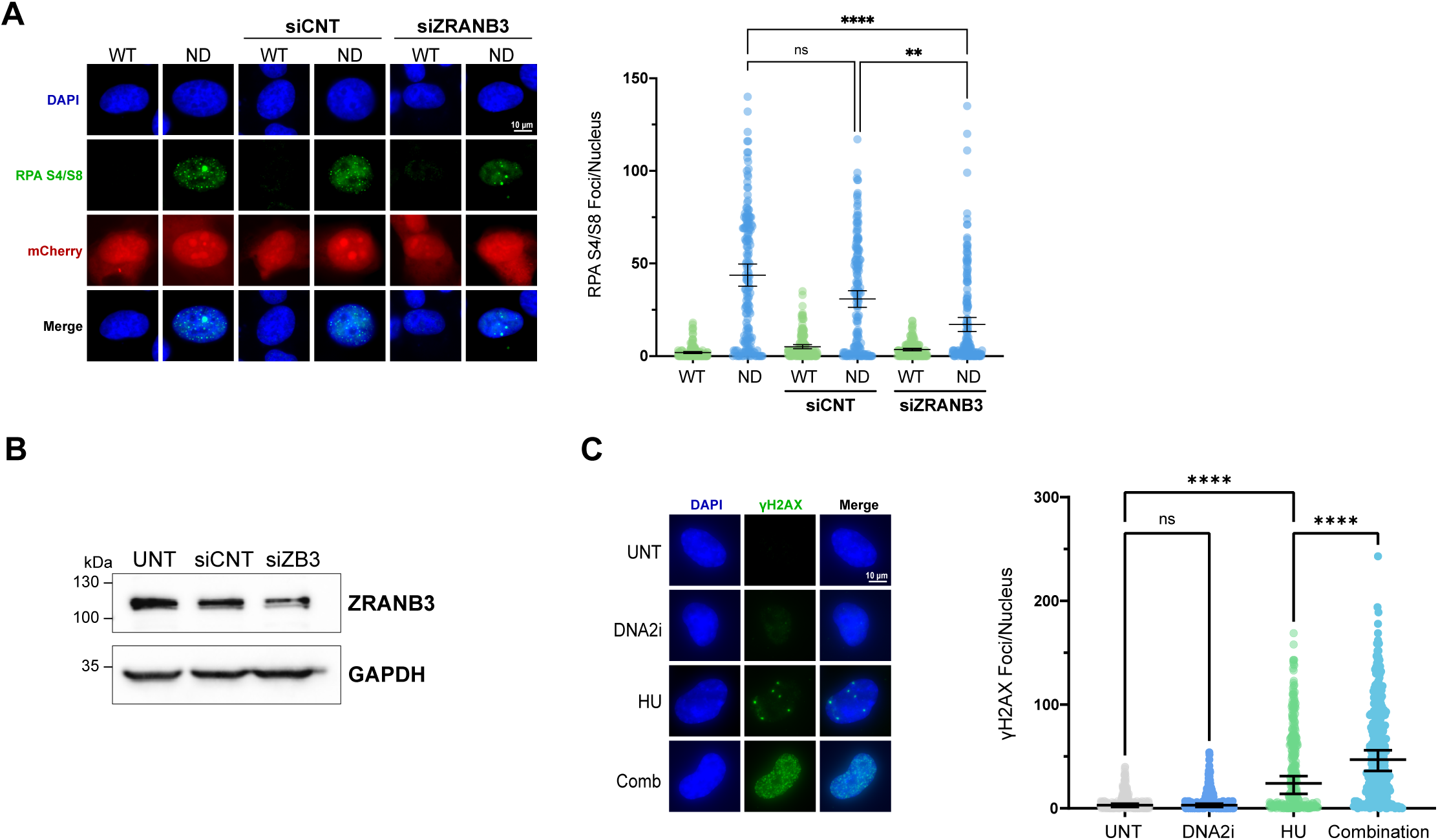
Replication fork reversal as a candidate driver of DNA2-nuclease inhibitor sensitivity. (A)Representative images (left) and quantification (right) of replication stress induction measured by phospho-RPA32-ser4/8 foci in cells with DNA2-WT or ND and the indicated siRNA. Quantification of foci numbers from A shows significant reductions in RPA S4/8 foci after ZRANB3 depletion. N=3; **P < 0.01, ****P < 0.0001 by Kruskal-Wallis test; mean ± 95% CI. (B) Representative western blot of ZRANB3 after siRNA depletion. (C) Representative images (left) and quantification (right) of γH2AX foci per nucleus in RPE1-hTERT cells exposed to a sub-lethal dose of DNA2 inhibitor in the presence or absence of hydroxyurea. ****p<0.0001 by Kruskal-Wallis test; mean ± 95% CI.

## DISCUSSION

There has been growing interest in DNA2 as an anti-cancer target owing to its roles in responding to replication stress in various contexts including oncogene-activation and ALT telomeres (Kumar et al, 2017; Ragupathi et al, 2025). However, targeting a protein with multiple catalytic domains and a variety of roles in genome maintenance presents a challenge as to how and when it should be targeted. Here we use separation-of-function mutants to show that expression of nuclease-dead alleles produces a dominant negative phenotype. This phenotype is dependent on helicase activity and DNA binding, raising interesting potential mechanisms. The role of DNA2’s helicase activity is incompletely understood. In biochemical studies the helicase activity of DNA2 is cryptic, revealed only in the nuclease dead mutant, which unwinds kilobases of DNA (Pinto et al, 2016). It has been suggested that this cryptic activity would be dysregulated in cells when DNA2 ND is expressed, leading to the observed accumulation of RPA-coated ssDNA and activation of the replication stress response.

The translocase activity of DNA2’s helicase domain has been shown to be important for strand specificity and processivity when digesting long ssDNA tracts (Zhou et al, 2015; Miller et al, 2017; Levikova et al, 2017). It acts directly against loading of a 3’ end, maintaining strand specificity. It was also shown that the ATPase dead mutant significantly slowed processing of long ssDNA flaps, especially when coated with RPA (Levikova et al, 2017). It is therefore possible that the translocase activity plays a role in loading of DNA2, assisting with RPA displacement. In this model, DNA2 ND completely loads onto the 5’flap and becomes trapped due to lack of cleavage activity and the inability to unthread in the 3’-5’ direction. The phenotype is therefore rescued in the ND/HD double mutant due to transient or weakened loading onto the substrate, preventing protein trapping. Both models are supported by our finding that disruption of RPA displacement through deletion of the α1-OB domain rescues the mutant phenotype. Fractionation experiments showing retention of DNA2 ND in the chromatin compartment favour a trapping mechanism. Protein trapping has been a successful strategy in clinical PARP inhibition (Mateo et al, 2019). Thus the possibility of replicating this mechanism in a DNA2 inhibitor is a promising area for future development.

Our results and others have implicated the ALT phenotype in sensitivity to DNA2 loss or inhibition. DNA2 has been shown to process BLM-generated long Okazaki flaps at telomeres (Jiang et al, 2024). Others have demonstrated an increase in TERRA R-loops upon DNA2 depletion in ALT cells (Ragupathi et al, 2025). DNA2 also has known roles in G-quadruplex (G4) processing at telomeres (Fernandez et al, 2025; Lin et al, 2013). With multiple potential substrates, it is perhaps unsurprising that multiple ALT cell lines were sensitive to DNA2 nuclease mutation or inhibition. However, our results show that ALT status alone is not a sufficient biomarker for sensitivity. In a panel of osteosarcoma cell lines, two out of the three tested ALT lines were not sensitive to nuclease inactivation. Furthermore, two out of three TERT lines were sensitive to the mutant. Nearly all of the ALT and TERT cell lines tested herein have multiple sources of replication stress with the potential to induce replication fork reversal. It is notable that the sensitive OS lines U2OS, SAOS2 and OS833 do not express full-length ATRX by western blot, while the other ALT lines are ATRX positive (Juhász et al, 2018). Given the specific effects of ATRX on telomeric DNA repair intermediates, it is possible that ATRX-negative ALT cells are a key feature of DNA2 nuclease sensitivity. Additional work is needed to unequivocally identify robust biomarkers to select patient populations most likely to benefit from DNA2 nuclease inhibition.

To gain a deeper understanding of the mechanism behind the DNA2 ND phenotype, we sought to understand which DNA2 substrate was involved. The siRNA knockdown of fork reversal factor ZRANB3 partially rescued the phenotype, showing that fork substrates may be key recruiters of the ND mutant. Furthermore, induction of replication fork reversal in RPE1 cells by hydroxyurea treatment strongly sensitized them to DNA2 nuclease inhibition. Our results align with a recent publication showing that processing of reversed replication forks underlies DNA2’s essentiality in yeast and human cells (Hudson et al, 2025). These authors further showed DNA2 depletion causes accumulation of RPA-coated ssDNA in G2 and subsequent senescence. It is possible that the arrest and growth restriction observed in cells expressing DNA2 ND is the result of a similar senescence phenotype. Future work to understand the dominant negative phenotype of DNA2 alleles and place them in the context of specific cancer genomic states will inform the development of strategies to inhibit DNA2 in targeted therapy.

## MATERIALS AND METHODS

### Cell culture conditions

The U2OS and Saos2 cell lines were cultured in McCoy’s 5a medium. HeLa, HeLa 1.3, HEK293T, and VA-13 were cultured in Dulbecco’s modified Eagle’s medium (DMEM). RPE1-hTERT cells were cultured in DMEM/Ham’s F12 50:50 medium. Media were supplemented with 10% fetal bovine serum and cultures were maintained at 37°C and 5% CO2.

### Cloning, plasmids, and mutagenesis

The p.CMV.attr.PGKmCherry expression vector was a Gibson assembly of pcDNA3.1 and pRRLSIN.cPPT.PGK-mCherry.WPRE which was a gift from the Andrew Weng lab. The flag-tagged expression vector for DNA2 was a generous gift from the Pavel Janscak lab (Sturzenegger et al, 2014). Mutations in DNA2 were introduced by site directed mutagenesis using the QuikChange Lightning kit (Agilent Technologies) following the manufacturer’s protocol. Where necessary, mutations were moved between vectors using PCR and restriction enzyme cloning. All plasmids were verified by full plasmid sequencing from Plasmidsaurus.

### Plasmid and siRNA transfection

Cells were transfected for 48-72 hours using Lipofectamine® 3000 according to the manufacturer’s protocol (Invitrogen). Cells were incubated for 48 – 72 hours post transfection prior to analysis. Transfected cells were identified by a fluorescent marker. For experiments involving siRNAs, cells were reverse transfected with siZRANB3 (Dharmacon 84083) using Lipofectamine® RNAiMAX reagent following the manufacturer’s protocol (Invitrogen). For co-transfection experiments, cells were incubated 24 hours prior to media change and forward plasmid transfection as described above.

### Immunofluorescence and analysis and statistical methods

For immunofluorescent (IF) analysis, cells were seeded on 8 well chamber slides overnight prior to transfection. For positive control treatments, cells were treated with either 3 µM mitomycin C or 10 µM cisplatin overnight. 48 hours post transfection, cells were fixed in 4% paraformaldehyde (PFA) in PBS for 15 minutes. Fixation was neutralized with 10 mM PBS glycine for 10 minutes followed by a PBS wash. Blocking and permeabilization were performed in saponin buffer (0.05% saponin, 0.2% BSA in 1X PBS) for 30 minutes followed by overnight treatment with primary antibodies. Antibodies were diluted as follows in saponin buffer: H2AX CST9718 Rabbit (1:1000) or Sigma 05-636 Mouse (1:500); RPA ab2175 (1:200); RPA Phosphoserine 4/8 Sigma PLA0071 (1:500); RPA Phosphoserine 33 Bethyl A300-246A (1:200); FLAG Sigma M2 F1804 (1:500). The next day, cells were washed 3X in saponin buffer followed by staining 1 hour in appropriate secondary antibodies: AlexaFluor® Rabbit 488 (1:1000); AlexaFluor® Rabbit 568 (1:1000); AlexaFluor® Mouse 568 (1:1000); AlexaFluor® Mouse 647 (1:500). Following secondary staining, cells were washed 3X in PBS and rinsed 3X in distilled water prior to sealing with Duolink In Situ Mounting Medium with DAPI.

For drug combination immunofluorescence experiments, cells were seeded at 0.75 × 10^6 cells and on day 1, HU was added for 4 hours, after which NSC-5195242 was added for an additional 4 hours in the combination group. Following drug treatment, cells were stained for DNA damage foci as previously described. For TeloFISH experiments, as described in Jiang et al., (2024), (145) after secondary antibody treatment and washing, cells were re-fixed in 4% PFA for 10 minutes at room temperature. Telomeres were then probed using the Dako Telomere PNA FISH Kit Cy3 (K5326) following manufacturer’s protocol.

All the imaging except for **Fig 3E and S4,** was performed using a Leica inverted DMi8 microscope at 65X or 100X magnification. Foci counting was performed in ImageJ using the find maxima tool.

For imaging using Image Xpress Confocal HT.ai (Molecular Devices), U2OS and RPE1 cells were seeded (30000 cells/well) on day 1 into a 96 well optical bottom black plate with a polystyrene film at the bottom (Nunc). Increasing concentrations of NSC-5195242 was added to both lines on day 2 followed by fixation and staining with ɣH2AX, RPA S33, and DAPI on day 3 and 4 as previously described. The plate was imaged with the Image Xpress Confocal HT.ai with a 20x WaterApo LambdaS WD 0.95 NA water immersion objective in confocal mode (50 μm slit).The images were captured and collected using MetaXpress 6.7.2.290 acquisition software, later post processed and analysed using IN Carta image analysis software with advanced AI.

For all microscopy experiments, the significance of the differences was determined using Prism version 5 or higher (Graphpad).

### Western blotting

Whole cell lysates were harvested in NP-40 buffer supplemented with benzonase nuclease, 1 mM Mg2+, protease inhibitor (Thermo-Fisher Scientific, A32955), and phosphatase inhibitor (Sigma, 4906845001). Protein concentration was estimated by Bio-Rad Protein Assay and equivalent amounts of protein by weight were resolved by SDS-PAGE. Proteins were transferred to a nitrocellulose membrane using a wet transfer and blocked with 3% milk in tris-buffered saline (TBS). Membranes were probed overnight at 4°C with the following primary antibodies diluted in 3% milk-TBS: FLAG M2 CST14793 Rabbit (1:500); Beta-5-tubulin ab15568 (1:5000); GAPDH GA1R (1:3000); Histone H3 ab1791 (1:20000). The next day, membranes were washed in TBS with 0.1% Tween-20 (TBS-T) and probed with anti-Rabbit IgG and anti-Mouse IgG antibodies conjugated to horseradish peroxidase (HRP). Following secondary staining, membranes were washed in TBS-T and developed using enhanced chemiluminescent HRP substrate.

### Chromatin fractionation

Fractionation was performed using a protocol adapted from Zhong et al. (2013). Cells were resuspended from 6 well plates, washed with PBS, and lysed in Low Salt Buffer + 1% Triton X-100 (10 mM Tris-HCl pH 7.4, 0.2 mM MgCl2) with protease inhibitor (Thermo-Fisher Scientific, A32955) and phosphatase inhibitor (Sigma, 4906845001). The supernatant was collected after pelleting at 14 000 rpm for 10 minutes at 4°C. The chromatin pellet was then washed 3 times in PBS and resuspended in 0.2 N HCl. After 20 minutes incubation on ice, the supernatant was collected and neutralized with an equal volume of 1 M Tris-HCl, pH 8.0. Fractions were analyzed by western blotting as described above with Ponceau S staining prior to blocking. Where applicable, western blots were analysed using Image Lab (Bio-Rad Laboratories) using the lanes and bands tool.

### DNA damage flow cytometry

Flow cytometric analysis of nuclear proteins was performed with the Transcription Factor Buffer Set from BD Pharmingen (#562574). Approximately 2 million cells per condition were fixed with the transcription factor fix buffer for 1 hour at room temperature. Cells were washed in perm/wash buffer and resuspended in primary antibodies: mouse anti-H2AX Sigma 05-636 (1:400) and rabbit anti-RPA Phosphoserine 4/8 Sigma PLA0071 (1:400) for 30 minutes at 37°C. All antibodies were diluted in perm/wash buffer. Cells were then washed and resuspended in secondary antibodies: AlexaFluor® Rabbit 488 (1:400) and AlexaFluor® Mouse 647 (1:400) for 30 minutes at 37°C. Following 2X washes in PBS, cells were resuspended in PBS with 0.5 µg/mL DAPI and incubated for 15 minutes at room temperature. Flow cytometric analysis was performed on the BD LSRFortessa analyser and the resulting data was analysed with FlowJo software.

### Incucyte cell growth assay

Cells were transfected in 6-well format as previously described and incubated for 5 hours at 37°C. Transfected cells were resuspended in complete growth medium and 30 000 cells/well were seeded in 96-well plate format, with 6 technical replicates per condition. Plates were imaged every 2 hours over a 5-day period for both brightfield and mCherry expression using an Incucyte SX5 live cell imager (Sartorius). Images were analysed to identify the number of mCherry+ cells per condition on the Sartorius Live-Cell Imaging and Analysis Software.

### ATP Glo viability assays

For DNA2 mutant experiments, cells were transfected in 6-well format as previously described. Two days post transfection, mCherry positive cells were sorted into an opaque 96-well plate, 1000 cells/well, 8 technical replicates per condition. For the untreated control, 1000 live cells were sorted into each well. Cells were maintained for 5 days after sorting, at which point they were treated with the CellTitre-Glo® 2.0 reagent following the manufacturer’s protocol (Promega). Luminescence was measured on the Tecan Spark® plate reader. To monitor the effect of DNA2 inhibition, cells were treated with NSC-5195242 on day 1 at respective concentrations and followed by Glo assay on day 4.

### Colony formation assay

Prior to seeding, cells were cultured as previously described. Drugs were prepared in 6-well plates in serial dilutions at 2x the desired final concentration. An equal volume of cells was added to each well at a low 500-2000 cell/well seeding density. Cells were incubated for 72 hours in drug, followed by a media change and outgrowth for 7-10 days. When colonies had reached sufficient size for counting, cells were washed and fixed in 4% PFA in PBS for 15 minutes at room temperature. Colonies were stained in crystal violet (Millipore Sigma #548-62-9) for 30 minutes prior to washing and quantification. Colonies were quantified by counting or total area using the “Colony Area” plugin in ImageJ and normalized to the untreated condition.

### DNA2 CETSA

Cells were treated with 10 µM NSC5195242 and harvested in NP-40 lysis buffer supplemented with benzonase nuclease, EDTA-free protease inhibitor cocktail, and phosphatase inhibitor. Lysates were clarified by centrifugation, and equal amounts of protein were aliquoted into eight fractions. The fractions were then subjected to a thermal gradient (0–58 °C) using a PCR thermal cycler to induce temperature-dependent protein denaturation. Following heating, the soluble supernatants were collected, and Western blotting was performed as described. DNA2 18727-1-AP Rabbit (1:600) and GAPDH (loading control) were used as primary antibodies.

## ACKNOWLEDGEMENTS

This work was supported by a Terry Fox Research Institute program project grant (#1155) to P.C.S. and a Canadian Institutes of Health Research scholarship to K.E.B. Plasmid reagents were generously provided by the Andrew Weng and Pavel Janscak labs. Cell lines were provided by Alejandro Sweet-Cordero. We thank members of the Stirling lab for helpful discussions.

## Notes

### Competing Interest Statement

The authors have declared no competing interest.

### Summary of Updates

A new panel showing representative images of cells in Figure 5C was added along with associated figure legend changes.

