## Supplementary Information File for "DNA2 variant analysis supports the nuclease activity as a preferred therapeutic target"

### SUPPLEMENTAL FIGURE LEGENDS

**Supplemental Figure 1.** Expression of nuclease dead DNA2 mutants produces a dominant DNA damage phenotype. (a) Schematic of the DNA2 gene. Mutations highlighted by arrows. Representative images (b) and quantification (c) of γH2AX foci in U2OS cells 48 hours post transfection with the indicated DNA2 mutants. Untransfected cells treated with 3 μM MMC were used as a positive control. UNT = untreated, EV = empty vector, MMC = mitomycin C. N=3; ****$P < 0.0001$ by Kruskal-Wallis test; mean ± 95% CI.

**Supplemental Figure 2.** (a) Western blot showing expression of wildtype and mutant DNA2 in U2OS cells; N=3. (b) Same as in A with K300R/K654E mutant; N=2. ****$P < 0.0001$ by Kruskal-Wallis test; mean ± 95% CI.

**Supplemental Figure 3.** Measuring the phenotype of DNA2 N-terminal deletion mutants by flow cytometry. Quantification of (a) γH2AX and (b) RPA phosphoserine S4/S8 positive cells detected by flow cytometry. U2OS cells were transfected with the indicated plasmids for 48 hours prior to staining and flow cytometric analysis. N=3; mean ± SD. Indicated P-values by One-way ANOVA.

**Supplemental Figure 4.** (a) Representative images of γH2AX and RPA phosphoserine 33 staining of RPE1-hTERT or U2OS cells treated with the DNA2 inhibitor NSC-5195242 in increasing concentrations. (b)The C5 global DNA2 inhibitor is more toxic to the RPE1 cell line than U2OS**.** Relative survival of U2OS and RPE1 cells treated with the indicated concentrations of the DNA2 inhibitor C5. Cells were treated for 72 hours followed by approximately 7 days outgrowth. Colonies were fixed and stained with crystal violet prior to counting. Survival normalized to untreated well for each plate. Dose response curves plotted with nonlinear regression. N=3, mean ± SD.

**Supplemental Figure 5.** (a) ALT-associated PML body (APB) assay for verification of ALT status in osteosarcoma cell lines. A telo-FISH assay was performed followed by immunofluorescent staining of PML for detection of co-localization in the indicated cell lines. (b) Cellular Engagement Thermal Stability Assay (CETSA) results showing soluble DNA2 remaining as a function of temperature by western blot in the presence of DMSO or NSC-5195242.

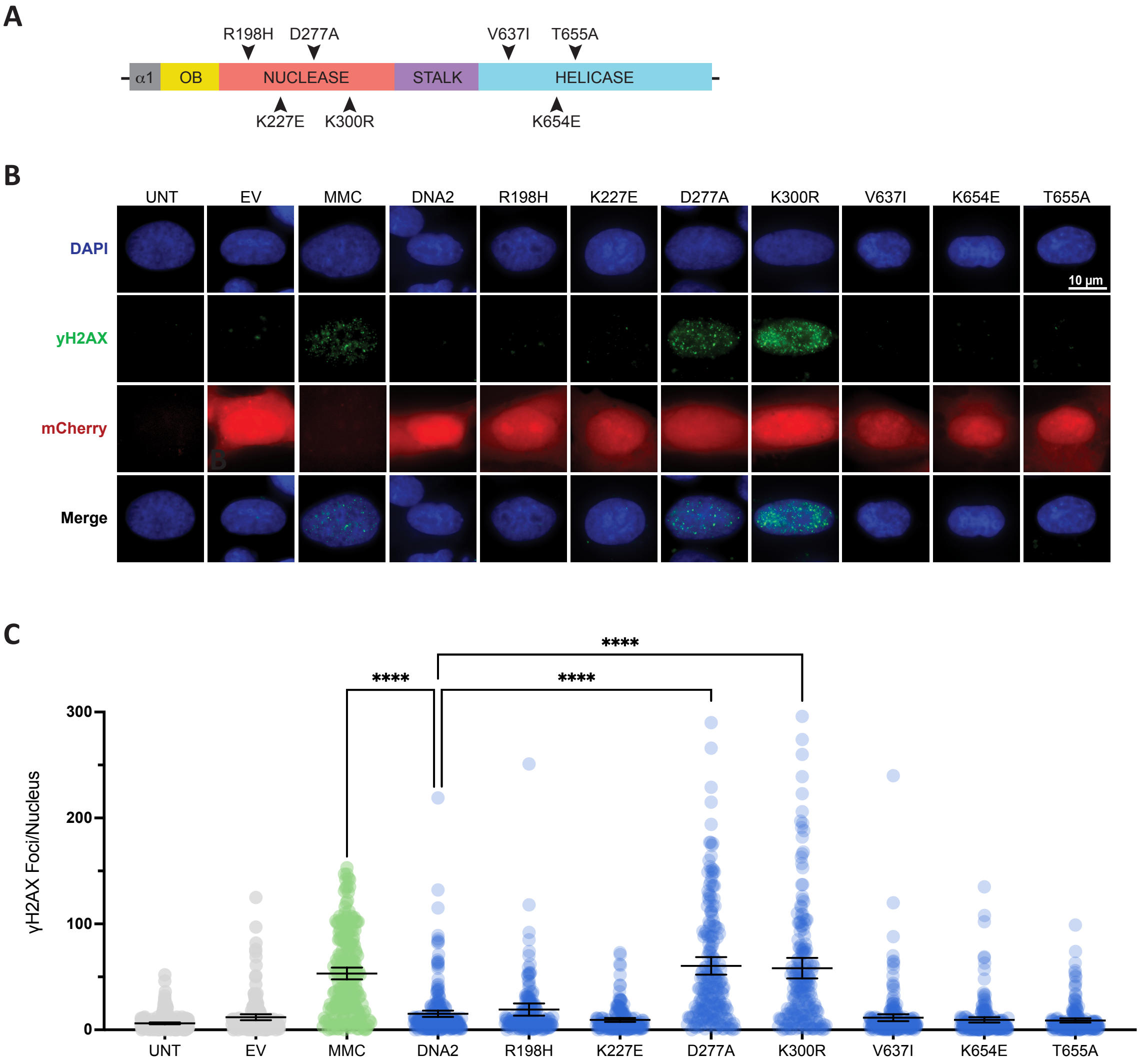


SUPPLEMENTARY FIGURE S1

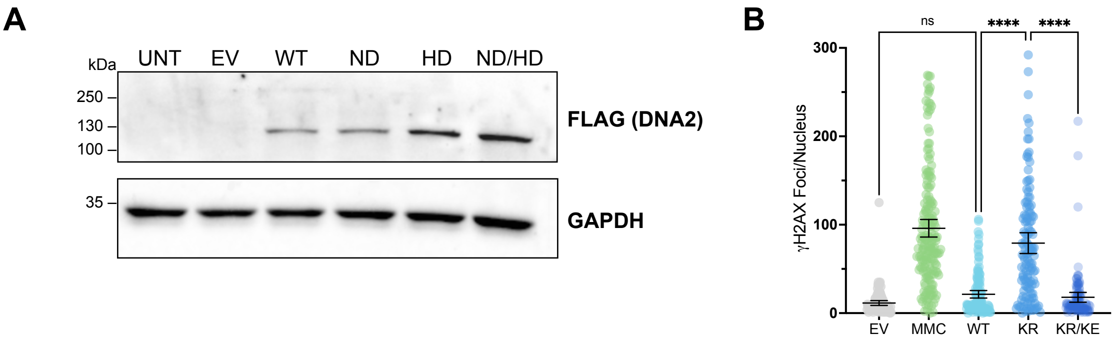


SUPPLEMENTARY FIGURE S2

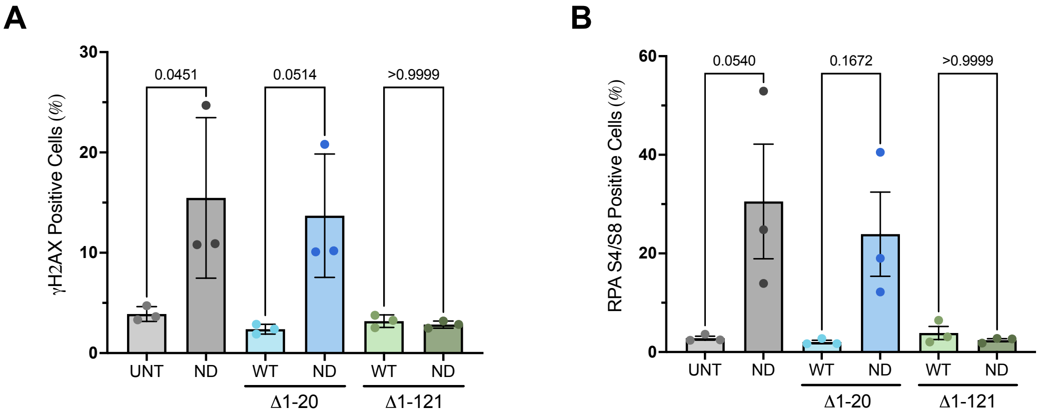


SUPPLEMENTARY FIGURE S3

A

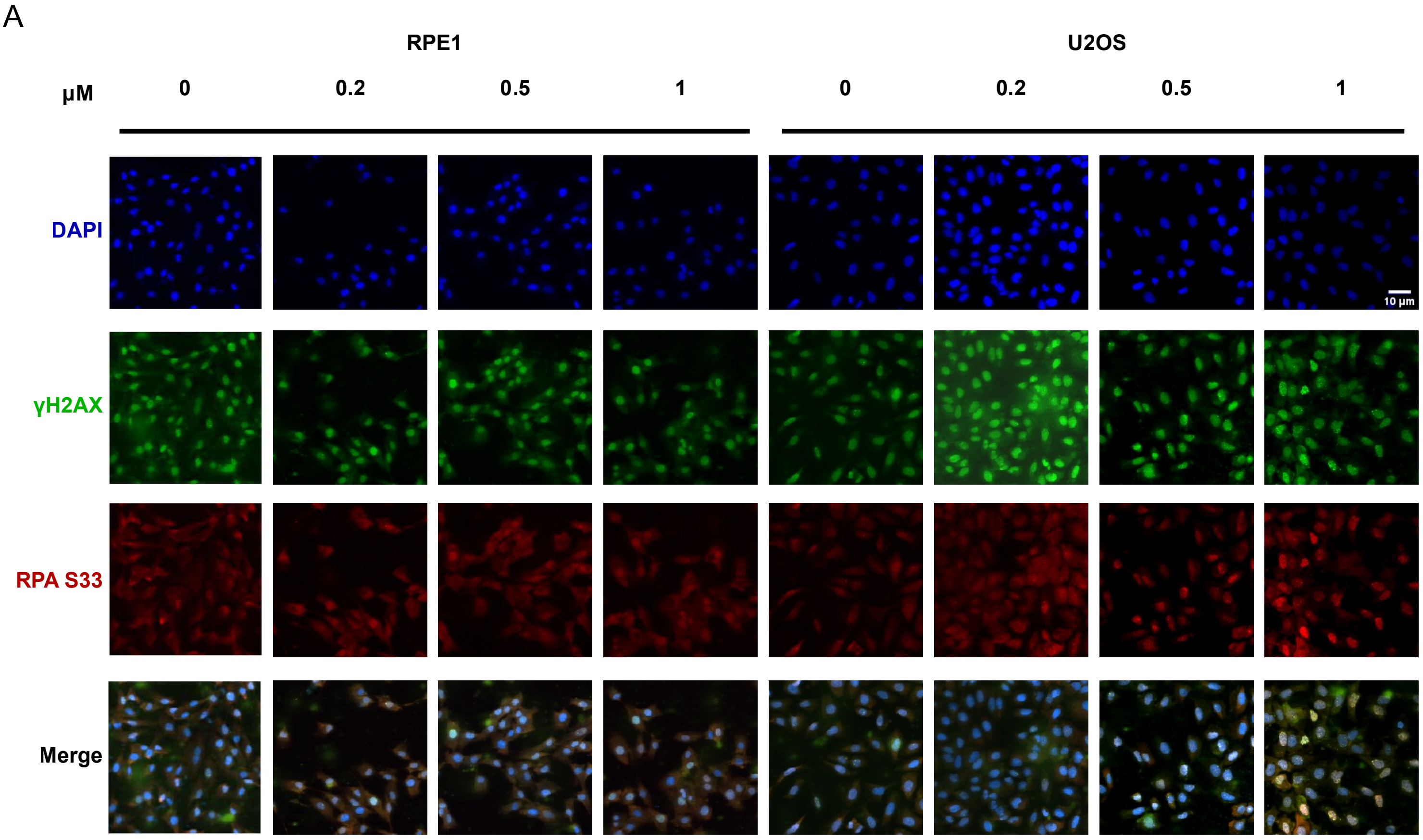


B

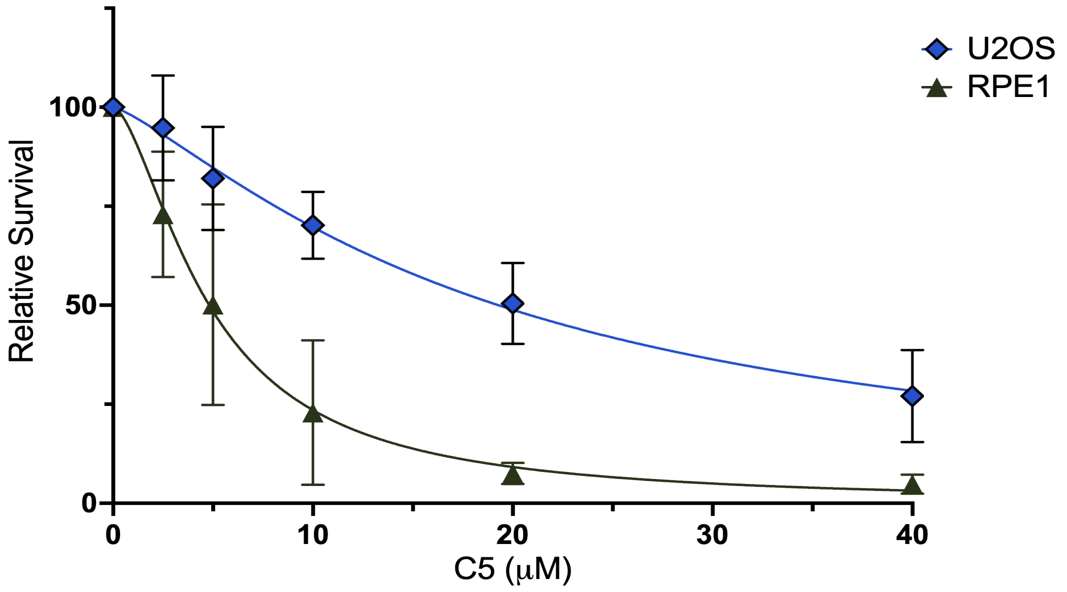


SUPPLEMENTARY FIGURE S4

A

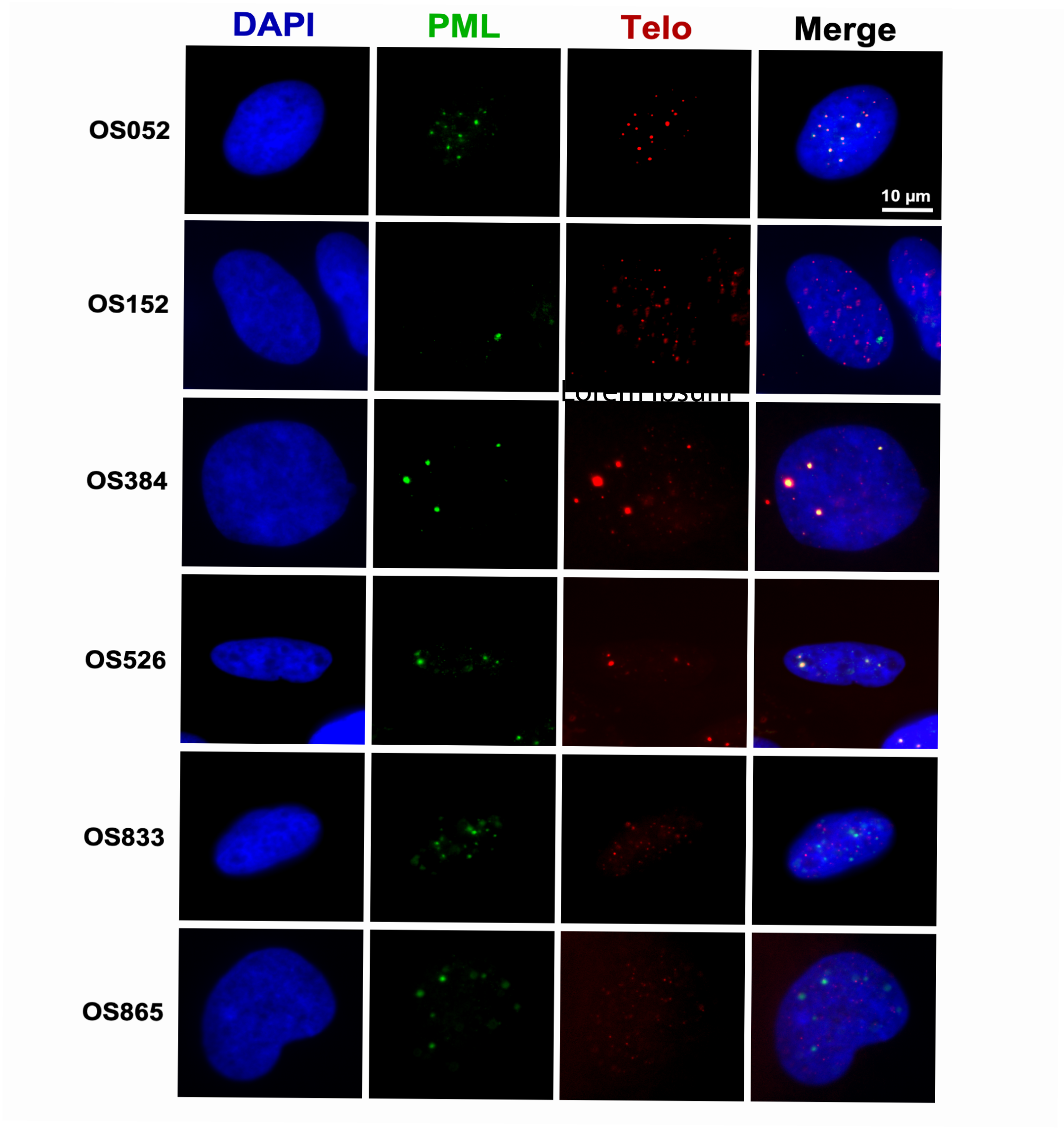
DAPI
PML
Telo
Merge
OS052
OS152
OS384
OS526
OS833
OS865
10 μm

B

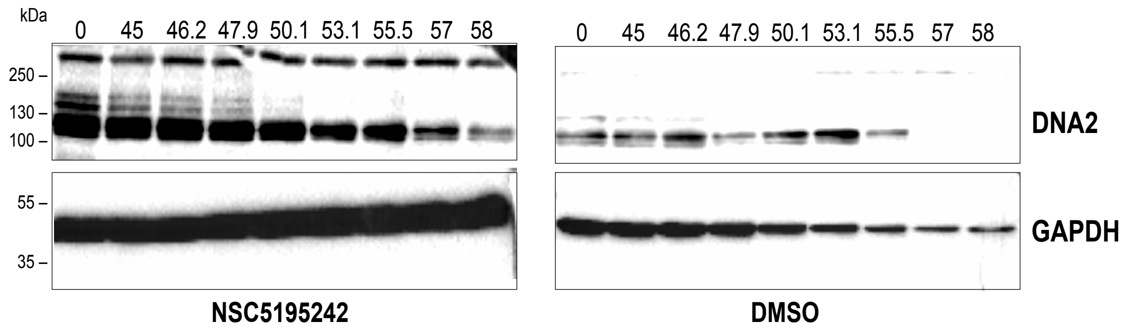
kDa
0
45
46.2
47.9
50.1
53.1
55.5
57
58
250
130
100
55
35
DNA2
GAPDH
NSC5195242
DMSO
